# Evolution of neurometabolic frugality in harvester ants

**DOI:** 10.64898/2026.01.30.702859

**Authors:** Craig D. Perl, Zachary N. Coto, Robert A. Johnson, Meredith G. Johnson, Leland C. Graber, James Haas, Jane S. Waters, James F.A. Traniello, Jon F. Harrison

**Affiliations:** School of Life Sciences, Arizona State University, 427 East Tyler Mall, Tempe, AZ 85281, USA; Department of Biology, Boston University, Boston, MA 02215, USA; Department of Entomology, Cornell University, 129 Garden Ave, Ithaca, NY 14853, USA; Department of Biology, Providence College, 1 Cunningham Square, Providence, RI 02918, USA

**Author notes:** Institute of Aquaculture, University of Stirling, Stirling, FK9 4LA, UK.

## Abstract

Despite the importance of neurometabolic costs in brain size evolution, quantitative data on brain metabolic rates are lacking. We measured *ex vivo* brain metabolic rates among species of the ant genus *Pogonomyrmex* to differentiate the roles of sociality and body size in brain evolution in a phylogenetic context. Worker body size and colony size (a proxy for social complexity) vary significantly among *Pogonomyrmex* species and were positively correlated. However, sociality was not a determinant of brain energetics. Worker body size strongly affected brain metabolism: 38% of resting metabolic rate was attributable to brain metabolism in species with the smallest workers, compared to 6% in species with the largest workers. More derived species had strikingly lower mass-specific brain metabolic costs, suggesting that increases in worker body size have selected for neurometabolic frugality through reductions in brain mass-specific metabolic rate. Additionally, smaller worker body sizes may require higher brain mass-specific energetic costs to achieve comparable performance by absolutely smaller brains. Our study shows that the social brain hypothesis does not explain patterns of brain size in *Pogonomyrmex*, but body size and evolutionary history strongly influence brain evolution in regard to both size and metabolic cost.

## INTRODUCTION

Understanding the selective forces that drive brain evolution and its macroevolutionary patterns remains one of the most intriguing unresolved problems in biology ^1–6^. Body size strongly affects brain size; smaller bodies have relatively larger brains (Haller’s Rule^7–10^), but the mechanisms linking body-size related patterns to brain morphology and physiology remain unclear^6,9,11^. In terms of social and ecological influences on brain evolution, theories hypothesize that the behavioural and cognitive demands of group living^12,13^ or diet and its influences on foraging^14,15^ are major selective factors of brain size and mosaicism. Collaterally, the expensive tissue hypothesis^16^ emphasizes the unusually high metabolic expense of the brain and posits that trade-offs between costly neural and digestive tissue permitted the evolution of larger brain sizes. Furthermore, an adipose-brain trade-off may also explain brain size evolution, with fat stores and encephalization forming alternative insurance policies against starvation^17^. Support for these theories varies and debate continues^17–19^, most often focusing on vertebrates^1,20^, particularly humans and other primates^21^ (and references therein). While energy costs are important components of many of these theories, few studies have measured brain metabolic rates (MR), and no studies have examined directional evolution of the size and MR of brains, and how these interact with body size or social evolution.

There is considerable evidence that larger animals have absolutely larger but relatively smaller brains, and some evidence that brains of larger animals use less energy per gram. Brain size scales hypometrically in mammals^8,22,23^, birds^23–25^, reptiles^26^, amphibians^27^, fish^28^, arachnids^29^, and insects^30,31^, including bees^32–34^ and ants^35,36^. Brain MR has also been shown to scale hypometrically in mammals and insects^11,37,38^. The causes of the hypometric scaling of brain size and metabolism are hypothesized to be related to constraints on wholebody MR^8^ and/or selection on life history parameters, such as the need for smaller animals to maintain a behavioural capacity comparable to that of larger animals^39–41^. There may also be selection for energy efficiency and a longer lifespan in larger animals^39,41,42^. Larger brains may be more cognitively capable^43–45^, but body and brain size effects on behavioural capacities are inconsistent. Generally, small-bodied species do not have limited behavioural repertoires compared to larger-bodied species^39,46^, and the need for smaller individuals and species to compete, behaviorally and cognitively, with larger competitors^41^ may explain hypometric brain scaling. The brains of the smallest insects displace somata and lose nuceli to preserve axon number, increasing relative brain mass allocated to energetically costly neurons, which may generate higher brain mass-specific metabolic rate (MSMR) in smaller individuals^47–50^. However, no study has examined the scaling of brain MR in a phylogenetic context, therefore there are no data on the extent to which variation in brain metabolism is associated with relatedness, independent of body and brain size variation, or whether there are directional evolutionary tendencies in neurometabolism.

Eusocial insects have emerged as important models for analyses of brain evolution in animal societies due to their diversity and size variation^31,36,51–55^, and here we test for effects of body size, social complexity, and phylogeny on brain size and MR in harvester ants, *Pogonomyrmex*. The application of the social brain hypothesis to eusocial insects is controversial^55–57^. Worker sterility within eusocial species precludes the evolution of individual strategies of reproductive competition, demanding coalition formation and socio-cognitive skills, which require increased brain tissue and a high processing capability. Eusocial insect workers and groups exhibit individual and collective cognition, respectively^58^, but worker cognitive abilities do not usually directly enhance worker fitness. In eusocial species with larger colony sizes, a proxy for social complexity due to increased social interactions^35,52,59–62^, individual behavioural/cognitive load may decrease due to task specialisation and division of labour^61^. This could result in smaller, less energy intensive brains associated with increasing social complexity^51,63^. Here, empirical data are equivocal^21,55^. Comparisons of the evolutionary neurobiology of ants suggest that striking differences in colony size and degree of social complexity influence brain size and its energetic expense^64^. In some ants, species that form larger colonies have evolved workers with relatively larger brains^35^. Sociality is also associated with larger brains in some bees^33,34,65,66^ but the broader pattern in this clade also indicates a significant role for diet and life history^32,67^. Conversely, in monomorphic fungus-growing ants^68^ and vespid wasps^52^, increased social complexity is associated, respectively, with decreased relative size of brains and mushroom bodies, the brain compartments specialized for learning and memory^67^. Together these results suggest that the effects of social complexity on social insect brains are complex and likely affected by ecology, colony size, social system, and eusocial insect clade^53,56^.

Phylogenetic history also appears to play important roles in brain evolution. Different clades often have different absolute and relative brain sizes^6,9,32,36^, and can have different relationships between brain size and body size^6^. The Marsh-Lartet rule, which suggests an evolutionary trend of increasing relative brain mass in mammals, was recently supported for three mammalian orders including primates^6,69^. In bees, phylogenetic analysis revealed that brain size evolution is linked to evolutionary changes in voltinism and host specialization^32^. Understanding the directional evolution of brain size and MR, and how tightly these are constrained by body size, are important questions for evolutionary biology. Unfortunately, we currently lack studies of directional evolution and phylogenetic patterns for brain metabolism in any animal clade.

To address major gaps in our understanding of the evolution of neurometabolism, brain size and social system, we measured brain masses, whole-body MRs, and intact brain MRs *ex vivo* from workers of eleven species of seed-harvesting ants, *Pogonomyrmex*^70^. Ants in general^71^, and *Pogonomyrmex* in particular, provide an excellent model to address questions at the intersection of neurobiology, socioecology, and metabolism. A larger body size is known to be associated with a relatively smaller brain within leafcutter ant subcastes^36^, within *Cataglyphis* species^35^, across 70 species of ants^36^, and more generally in insects^9^. *Pogonomyrmex* have significant variation in body mass, colony size, and foraging ecology. The species studied here are all primarily granivorous and reproduce annually, enabling tests for effects of body size, colony size and phylogeny, without obvious confounding variables of life history or diet (Table 1). All 11 species inhabit grasslands of the southwestern United States and robust phylogenies of the genus exist^72^, along with assessments of colony size (Table 1). We determined the effects of body size and social complexity on brain and body MRs in a phylogenetic context to gain insight into possible evolutionary trends. We ask: how is the evolution of *Pogonomyrmex* linked with changes in body size, brain size, brain and whole-body MRs, and social complexity?

**Table 1:** Means of variables from workers of each species.

| Species | Body mass (mg) | Brain mass (mg) | Body metabolic rate (μWatts) | Brain metabolic rate (μWatts) | Worker number per colony | Colony size reference |
| --- | --- | --- | --- | --- | --- | --- |
| *P. apache* | 12.35 | 0.15 | 16.09 | 1.65 | 80 | Cole (1954)<sup>131</sup> |
| *P. badius* | 14.12 | 0.14 | 11.52 | 1.72 | 5000 | Tschinkel (2017)<sup>132</sup> |
| *P. barbatus* | 14.95 | 0.19 | 19.74 | 1.68 | 12000 | Gordon (1992)<sup>133</sup> |
| *P. californicus* | 6.08 | 0.11 | 6.75 | 1.60 | 3250 | RA Johnson, pers. obs. |
| *P. desertorum* | 6.03 | 0.10 | 5.66 | 1.64 | 500 | Creighton (1956)<sup>134</sup> |
| *P. huachucanus* | 4.29 | 0.08 | 4.11 | 1.60 | 150 | Creighton (1952)<sup>135</sup>; RA Johnson, pers. obs. |
| *P. imberbicus* | 2.12 | 0.06 | 4.91 | 1.45 | 125 | Heinze, et al. (1992)<sup>136</sup> |
| *P. maricopa* | 9.80 | 0.12 | 5.86 | 1.63 | 750 | RA Johnson, pers. obs. |
| *P. occidentalis* | 7.91 | 0.11 | 9.20 | 1.76 | 3880 | Lavigne (1969)<sup>137</sup> |
| *P. pima* | 1.68 | 0.05 | 5.02 | 1.41 | 250 | Johnson, et. al (2007)<sup>138</sup> |
| *P. rugosus* | 15.87 | 0.29 | 29.81 | 1.64 | 8600 | MacKay (1981)<sup>139</sup> |

## RESULTS

### i. Phylogenetic signal

There was a significant phylogenetic signal for brain MSMR (Fig. 1; Pagel’s λ = 1.00, p < 0.01), body MSMR (Fig. 1; Pagel’s λ = 1.00, p < 0.01), brain MR (Fig. 1; Pagel’s λ = 0.98, p < 0.01), and colony size (Fig. 1; Pagel’s λ = 1.00, p < 0.01). For brain mass-specific MR and body MSMR there was a trend for more derived species to have lower rates (Fig. 1). In contrast, for brain MR and colony size, there was a trend for more derived species to have higher values (Fig. 1). There was no significant phylogenetic signal for body mass (Pagel’s λ = 0.55, p > 0.20), brain mass (Pagel’s λ = 0.65, p > 0.20) or whole-body MR (Pagel’s λ < 0.01, p = 1.00). As such, subsequent analyses using body mass, whole body MR, or brain mass were conducted without any phylogenetic correction, but any analysis of brain MR, mass-specific MR or colony size accounted for phylogeny.

**Figure 1:**
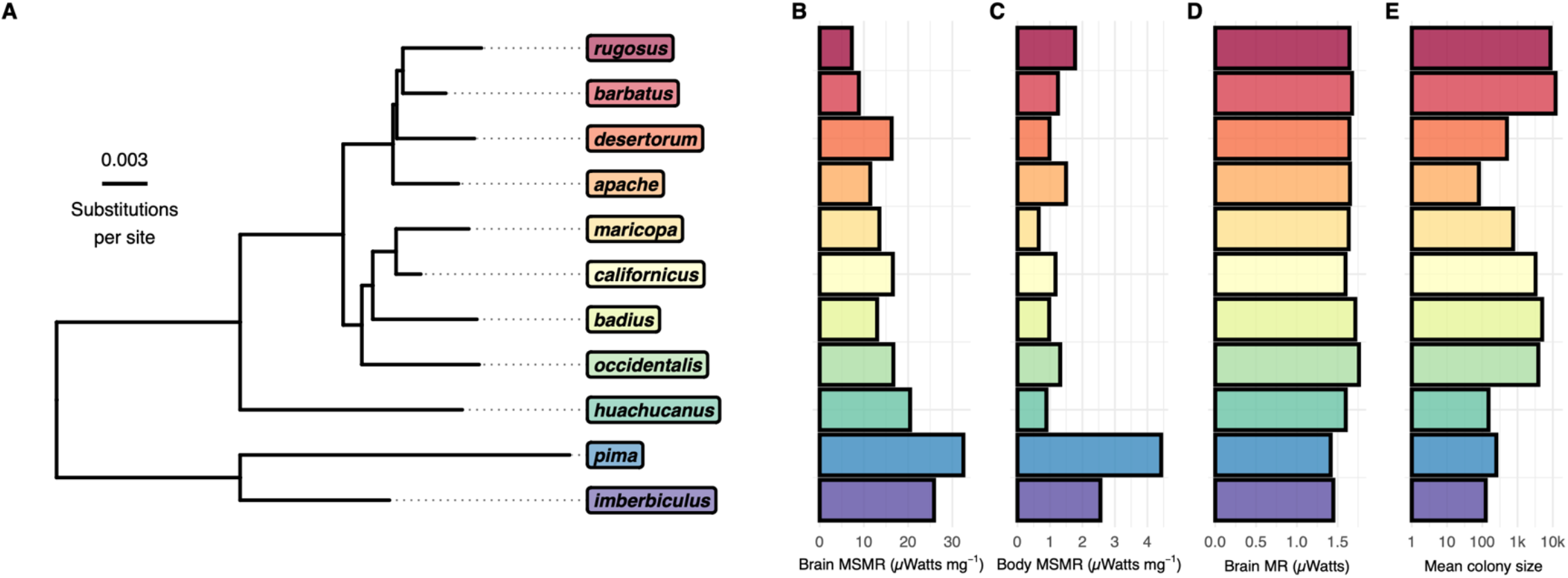
**(A)** Brain mass-specific and body mass-specific metabolic rates were strongly affected by phylogeny. **(B & C)** The basal clades of the genus had relatively high brain and body mass-specific metabolic rates, while the rugosus-barbatus clade had the lowest, suggestive of a progressive evolutionary trend toward lower brain and body mass-specific metabolic rates. **(D)** Absolute brain metabolic rates also had a significant phylogenetic signal; being generally lower in more basal species. **(E)** Mean colony sizes had a significant phylogenetic signal, being generally smaller in more basal species. Colours are not indicative of any value but included to aid readability.

### ii. Body size and colony size

Mean worker mass increased with increasing colony size (Fig. 1; PGLS; t_9,11_ = 2.73, p < 0.03).

### iii. Brain size scaling and relationship with colony size

There was a significant relationship between brain mass and body mass (Fig. 3A; linear mixed effects model; t_9,252_ = 15.73, p < 0.001). Brain mass scaled hypometrically, with an allometric slope of 0.54 ± 0.03 (for this and subsequent regression equations, the standard error is provided). There was a significant relationship between brain mass and colony size (Fig. 3B; PGLS; t_9,11_ = 3.07, p < 0.02): worker brain mass increased with increasing colony size. There was no significant relationship between relative brain mass (brain mass per unit body mass) and colony size (Fig. 3C; PGLS; t_9,11_ = 0.52, p = 0.62).

**Figure 2:**
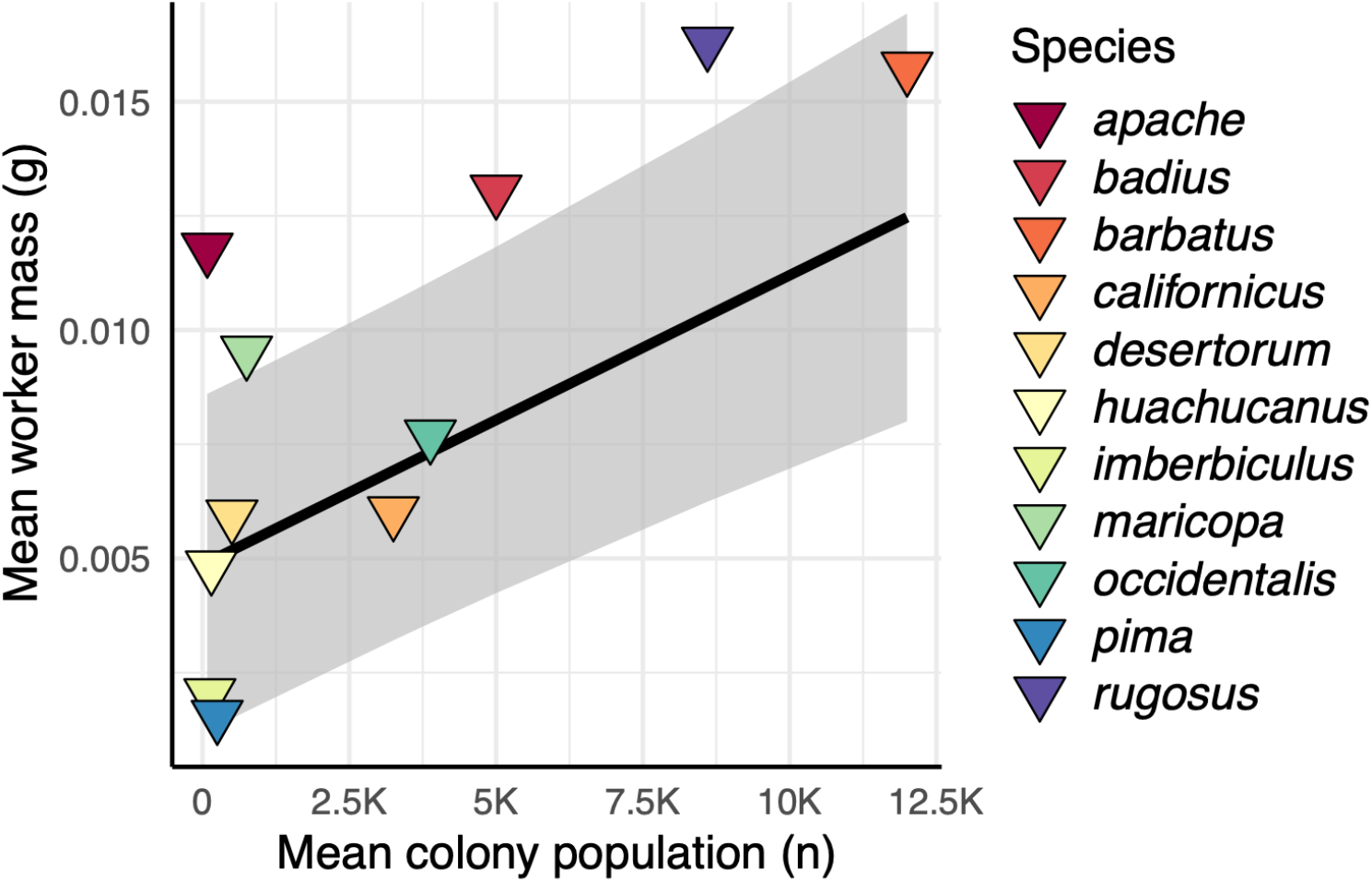
Worker masses increase with increasing social complexity. Mean worker mass increases with increasing colony size (worker mass = 0.000000634* colony population + 0.004862386).

**Figure 3:**
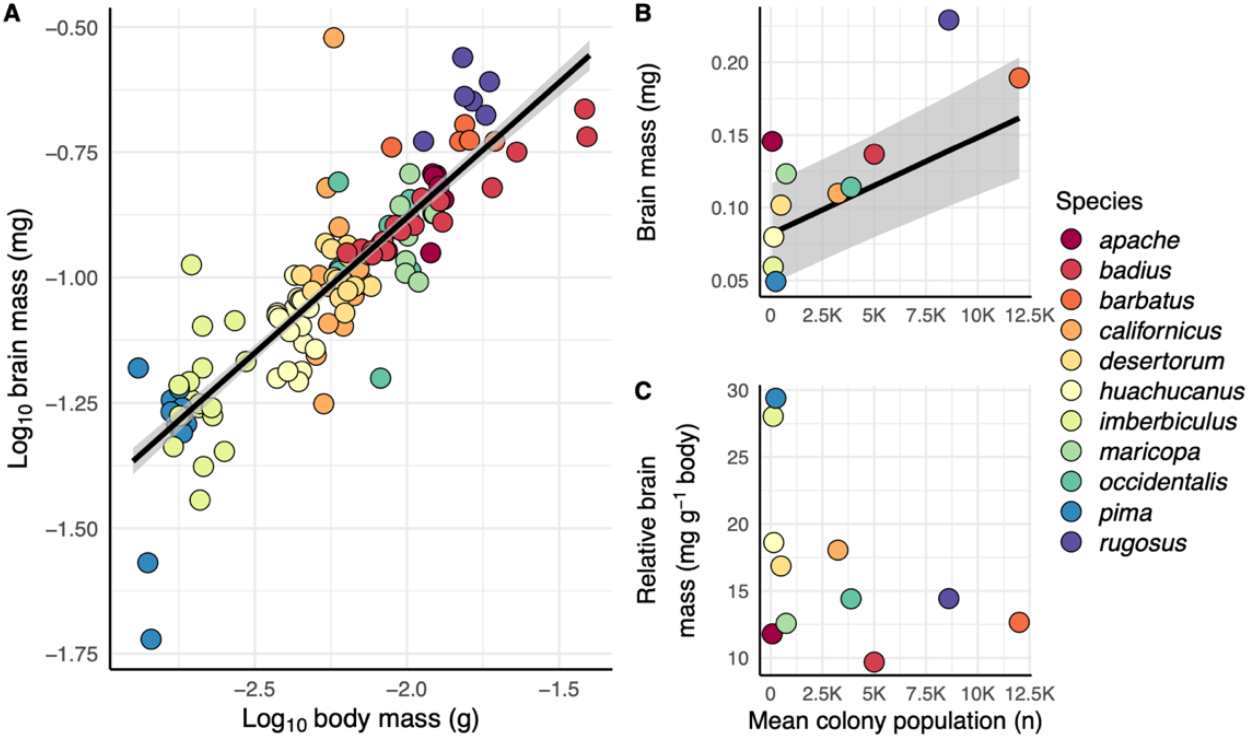
**(A)** Brain mass scaled hypometrically with worker body mass (log_10_ brain mass = 0.54 * log_10_ body mass +0.20) **(B)** Brain mass increased with increasing colonial population (brain mass = 0.00000666 * colony size + 0.08179035). **(C)** Relative brain mass was not significantly related to colonial population size.

### iv. Whole body resting metabolic scaling and relationship with colony size

There was a significant relationship between log_10_ whole body MR and log_10_ body mass (Fig 4A; linear mixed effects model; t_37,252_ = 3.10, p < 0.01). Whole body resting MR scaled hypometrically; the allometric slope was 0.35 ± 0.11. There was a significant relationship between whole body MR and colony size (Fig. 4B; PGLS; t_9,11_ = 2.44, p < 0.04). Whole body MR increases with increasing colony size. However, the relationship between whole body MR and colony size becomes nonsignificant if body size is taken into account. When colony size and body mass were both included as predictors of body MR in a multivariate model, body mass emerged as a significant factor (PGLS; t_1,11_ = 4.09, p < 0.01), and colony size does not (PGLS; t_1,11_ = 0.95, p = 0.37). There was no significant relationship between whole body MSMR and colony size (Fig. 4C; PGLS; t_9,11_ = 0.35, p = 0.73). Furthermore, there was no significant relationship between body-mass-corrected MR (residuals of whole-body MR regressed against body mass) and colony size (Fig. S2A; PGLS; F_1,11_ = 1.13, p = 0.29).

**Figure 4:**
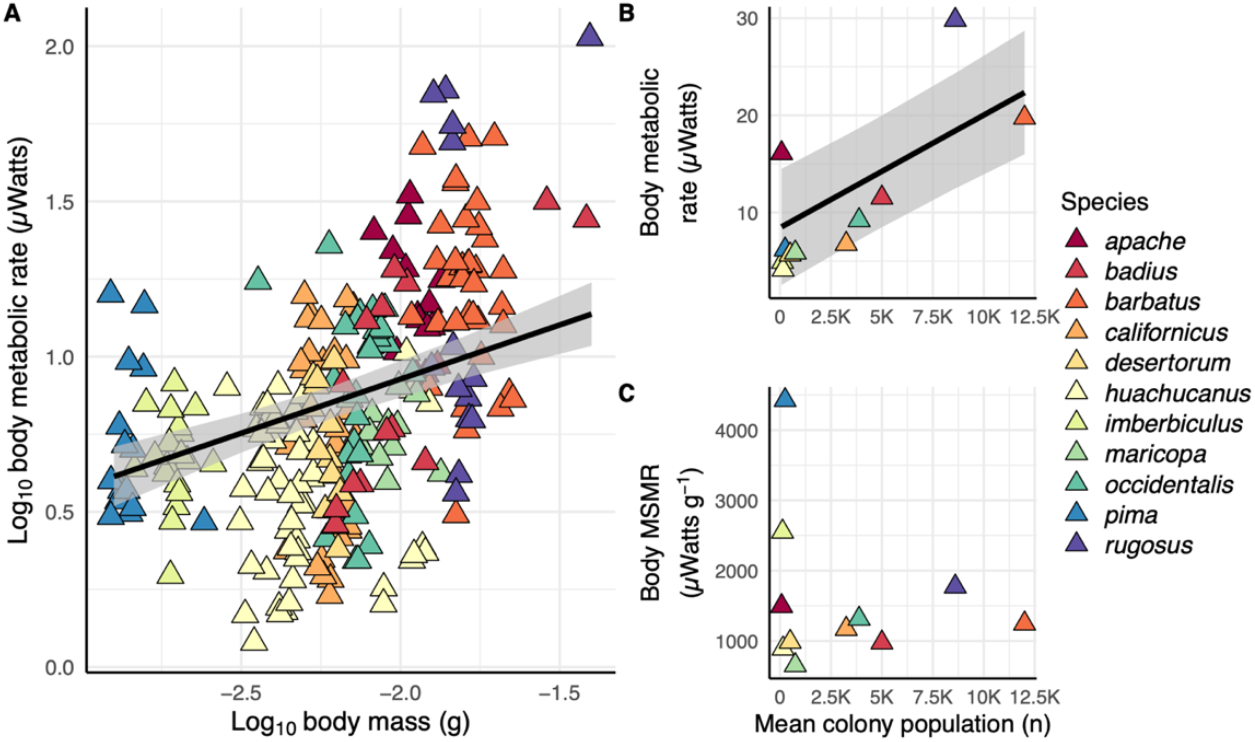
**(A)** Whole body metabolic rate increased in larger workers but scaled hypometrically (log_10_ body metabolic rate = 0.35 * log_10_ body mass + 1.62). **(B)** Whole body metabolic rate increased with increasing colonial population (Body MR = 0.001035 * colony size + 6.67). **(C)** Whole body mass-specific metabolic rate was not significantly related to colonial population size.

### v. Brain metabolic rate scaling and relationship with colony size

There was no significant relationship between brain MR and colony size (Fig. 5A; PGLS; t_9,11_ = 0.77, p = 0.46) and there was no significant linear relationship between brain MR and brain mass (PGLS; t_9,11_ = 0.84, p = 0.42). There was, however, a significant non-linear relationship between brain MR and brain mass (Fig. 5B; GAM with thin plate spline and phylogenetic penalty; F_1_,_11_ = 3.21, p < 0.01). Brain MR increased with increasing brain mass up to a value of 0.148 mg and then decreased slightly. There was a significant relationship between brain MSMR and colony size (Fig. 5C; GAM with thin plate spline and phylogenetic penalty; F_1_,_11_ = 7.06, p < 0.03). There was also a significant linear fit between brain MR and colony size (PGLS; t_9,11_ = 2.33, p < 0.05), but the non-linear model was selected due to having a lower AIC score (GAM = 70.89, PGLS = 75.02). Brain MSMR decreased with increasing colony size. There was also a highly significant relationship between brain MSMR and body mass (Fig. 5D; PGLS; t_1,11_ = 5.70, p < 0.001). As body mass increased, the relative energetic cost per gram of brain decreased; brains from smaller individuals used more energy per unit mass of brain than larger individuals. When colony size and body mass were both included as predictors of brain MSMR in a multivariate model, body mass emerged as a significant factor (PGLS; t_1,11_ = 5.70, p < 0.001), and colony size does not (PGLS; t_1,11_ = 0.68, p = 0.52). There was no significant interaction term (PGLS; t_1,11_ = 0.90, p = 0.40). Furthermore, there was no significant relationship between body-mass-corrected brain MSMR (residuals of brain MSMR regressed against body mass) and colony size (Fig. S2B; PGLS; F_1,11_ = 0.33, p = 0.58).

**Figure 5:**
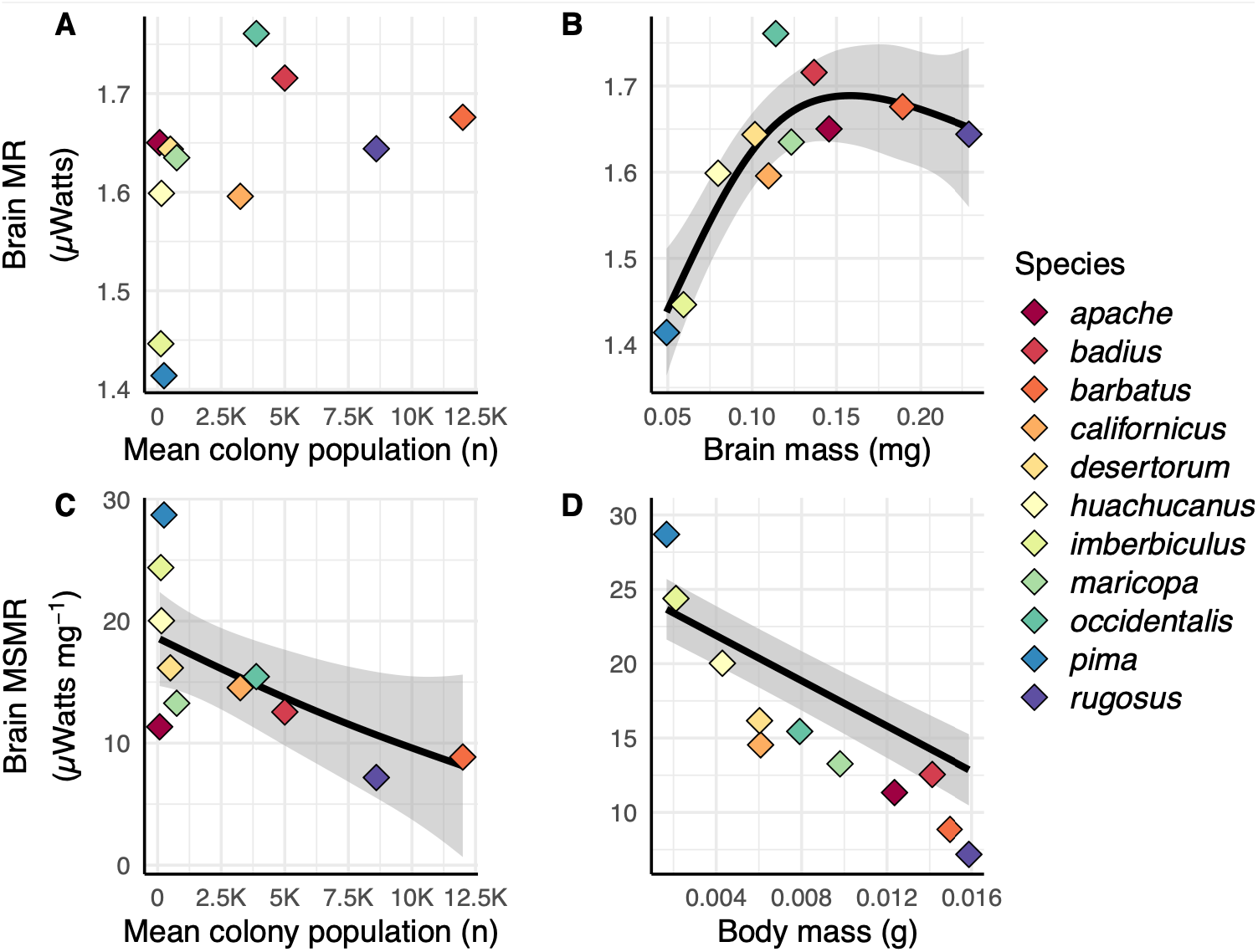
Phylogenetic generalized least squares analyses of independent effects on brain metabolism. **(A)** There was no relationship between brain metabolic rate and colonial population. **(B)** There is a significant relationship between brain metabolic rate and brain mass. **(C)** Brain mass-specific metabolic rate decreased with increasing colonial. **(D)** Brain mass-specific metabolic rate also declined with increasing worker body mass (Brain MSMR = -874.87 * body mass + 26.94). When analysing body mass and colony size as predictors of brain MSMR in a multivariate model, colony size ceases to be a significant predictor of brain MSRS and body mass remaining a significant factor.

## DISCUSSION

Phylogeny and body size had strong effects on brain and body energetics, but colony size, a proxy for social complexity, did not. There was a general trend for more derived species to have lower brain and body MSMR (Fig. 1). Body size did not have a phylogenetic signature, but was strongly correlated with brain MSMR, with larger-bodied species having lower MSMRs (Fig. 5D). Higher brain MSMRs were also correlated with increasing social complexity, but this relationship was non-significant when body size was accounted for (Fig. S2B). Brain MR increased with brain size only up to a brain mass of 0.148 mg (Fig. 5B). Species with larger brains (e.g. *P. barbatus, P. rugosus*) had equivalent brain MRs to those with significantly smaller brains (e.g. *P. apache, P. badius, P. occidentalis*). Colony size was strongly correlated with body size; larger colonies had larger workers (Fig. 1). There were some significant relationships between colony size and energetics: workers from larger colonies showed higher body MRs. However, this relationship was not independent of an increase in worker body size (Fig. S2A), and colony size was not a significant predictor of body MR with body mass accounted for. Therefore, body size was the more important factor for determining body and brain energetics. We identified strong directional evolutionary tendencies for these traits, with brain and body MSMR being lower in more derived species, indicating a directional evolutionary trend towards energetic frugality.

Though body size is the most important factor for determining energetics, how this trait affects the association of larger workers with larger colonies remains unclear. Worker size is inconsistently associated with colony size in social insects; some ant species (e.g. *Solenopsis invicta, Linepithema humile*) with the most populous colonies have relatively small workers and others are exceptionally polymorphic. However, across more than 100 ant species, worker mass, queen mass and colony mass are strongly positively correlated^74^. The positive association between worker size and colony size in our sampled *Pogonomyrmex* is therefore similar to broad, but not universal, trends across ants. The relationship between worker body size, foraging productivity, and energy efficiency is important but unclear, and may not reflect vertebrate patterns due to the impacts of larger colony sizes on worker resource acquisition. Across multiple vertebrate taxa, species with larger body sizes use more resources per individual, but as individual abundances and generation times scale inversely to individual size in vertebrates, large- and small-bodied species tend to utilize similar amounts of energy per year (the equal fitness paradigm^73,74^). There appear to be advantages to larger colony sizes. Larger colonies, associated with larger body-sizes in *Pogonomyrmex*, may control clumped resources^75,76^ (though see^77^) and avoid conflict with neighbouring colonies^78^, but it is unknown how resource acquisition compares across small and large colonies and whether they also conform to the equal fitness paradigm. Within *Pogonomyrmex*, group or trunk-trail foraging is restricted to larger bodied species and larger colonies^79^, therefore it is possible that the energetic benefits brought by the evolution of larger workers with access to differential foraging strategies can support larger colony sizes, though in other seed-harvesting species smaller individuals are associated with group foraging strategies^79^. It is therefore more likely that group foraging allows for resource domination in a typically stochastic environment. Individual foraging trips, irrespective of foraging strategy, have minimal impact on worker energy budgets^80,81^ but group foraging strategies can ease the exploitation of abundant resources, when they appear^79^. *Pogonomyrmex* colonies can experience significant interspecific food resource competition^76^ thus influencing the evolution of worker size through character displacement, reducing competition by niche segregation, or selecting for differential foraging strategy^82,83^. Colonies of *P. barbatus*, a species with the largest workers, have lower fitness when neighbouring colony density is higher^84^. Resource competition may favour the evolution of traits such as territoriality and trunk-trail foraging, enabling resource control and, collaterally, larger workers and colony sizes. Comparative data on foraging success, colony fitness, generation time, abundance, and fitness of these species in the field are lacking, and the hypothesis that large worker size, social complexity, and ecological dominance are linked in *Pogonomyrmex* requires additional research, especially to determine the causality among the evolution of colony size, body size, foraging strategy, and their metabolic consequences.

We did not find a significant effect of colony size on whole-body MSMR, perhaps suggesting that larger colonies are not more metabolically efficient in our sample of *Pogonomyrmex* species. However, it is important to note that colony MRs cannot be predicted from the resting MRs of individual workers^85^. Large colony sizes in social insects correlate with lower colony-level MSMRs^86^ as well as increased division of labour in *P. californicus*^61,87^. Worker task diversity increases as colonies grow^87^ with older, larger colonies having workers that tend to specialise on specific tasks, rather than engaging in a wider range of tasks^61,85^, in addition to lower activity rates^85,88^. Such behavioural specializations could allow larger colonies to reduce mass-specific energy use. There were extremely high effect sizes of phylogeny on brain MR, brain MSMR, and whole-body MSMR (Fig. 1). We did not find a phylogenetic signal for brain mass, diverging from evolutionary trends in mammals, where directional evolution for increasing relative brain mass has been reported within some mammalian orders, including primates^6,69^.

Although our analyses were performed on a single genus, the body size range across the *Pogonomyrmex* species in our study is comparable to the range exhibited within mammalian orders, allowing comparisons of scaling across these diverse clades. The distinct life histories and reproductive traits of mammals and eusocial insects may explain the differential neurometabolic response to increasing social complexity. Our data indicate that the social brain hypothesis may have limitations when applied to the brain energetics of eusocial insects^51^. We find concordance with other investigations into the association of sociality and neurometabolism in ants. Kamhi et al.^64^ report that Australasian weaver ants, *Oecophylla smaragdina*, a paradigm of insect social complexity, have larger brains and greater investment in the mushroom bodies but significantly lower cytochrome oxidase (COX) activity in the mushroom body medial and lateral calyces relative to those of the socially basic, small-colony sister species *Formica subsericea*. The mushroom bodies are a brain compartment dedicated to higher-order processing, integration, learning, and memory and thus likely provide neural support for complex social behaviour in ants ^63^. COX is a proxy for ATP usage; mass-specific brain energy expenditure may decrease as social complexity increases. Our data support this pattern by *directly* measuring brain energetics. We find the same relationship in *Pogonomyrmex:* brain MSMR decreases with increasing social complexity, rather than increasing as would be expected from social brain theory.

Phylogeny is more important for both brain and whole body MSMR, indicative of selection towards energetic frugality; the highest values occurred in the most basal clade (*P. imberbiculus* and *P. pima*), followed by the next most basal species (*P. huachucanus*). The lowest values occurred in the most derived clades, strongly suggesting an evolutionary trend toward lower brain and wholebody MSMRs. Absolute brain MR showed the opposite pattern: the lowest values were found in the more basal clades. Interestingly, brain and body mass showed no phylogenetic signal, suggesting that the above patterns are not simply driven by brain and body mass, and that mass is more evolutionarily labile than metabolic rate. Phylogenetic signal for body mass-related traits and MRs has been found to be minimal in some insects (e.g. stingless bees^89^, scarab beetles^90^), whereas others show strong effects (e.g. size-related variables of bees and moths^89,91,92^). Our sampling of 11 species in *Pogonomyrmex* should be extended to confirm these patterns, as we measured approximately a third of the known North American species^93^.

Larger-bodied ant workers have remarkably lower energy usage per unit mass of brain. The smallest-bodied species have 10x higher brain MSMR than the largest-bodied species in *Pogonomyrmex*. Due to their relatively large brains and higher brain MSMRs, brains account for a greater proportion of the metabolic budget in smaller-bodied species (38% in *P. huachuncanus*, 29% in *P. desertorum* and *P. imberbiculus*) but only 6% in the largest-bodied species (*P. rugosus)*. Even though brains account for only 2% of whole-body mass, this scaling of brain MR explains ∼25% of the hypometric scaling of whole-body MRs across the clade. This further demonstrates that body size is of high significance for determining neurometabolic costs and indicates that a small body size is associated with both relatively larger and more energetically active brains. The high energetic cost incurred by brains may provide a selective pressure to reduce relative brain MRs, as observed in the more derived *Pogonomyrmex* species.

The strong hypometric scaling of brain MR suggests either selection to reduce brain MRs in larger-bodied species, selection for high brain MRs in smaller-bodied species, or both. Limits to whole-body MR, such as increasingly constrained oxygen or nutrient delivery in larger brains, is a possible cause of hypometric brain MR scaling^8^. However, we currently lack studies of the scaling of oxygen and substrate supply in ants as have been performed for some other groups of insects^94,95^, and no studies have yet addressed this issue for any brains. Brains, and neural tissue in general, are often cited as being a significant constituent of overall energy budget^11,16^. As such, large-bodied species with the absolute largest brain masses might gain the most benefit from reducing brain MSMRs^11,96^. Larger-bodied species generally have longer lifespans (reviewed by^42^), and there is evidence that this is also true in ants^97,98^. Higher MSMRs are often associated with higher rates of reactive oxygen species production, which can damage tissues and shorten lifespan, therefore it is possible that the lower MSMRs of larger-bodied species arise ultimately from selection to reduce metabolic-linked tissue damage and extend lifespan^99,100^.

Alternatively, or additionally, species with smaller workers may experience greater selection to *increase* their brain MRs. Small harvester ant workers carry out the same tasks as larger-bodied species (e.g. forage for seeds, defend territory and/or food sources, navigate, build and maintain the nest, care for the queen and brood) with brains and sensory organs that are absolutely smaller; this may require increased intensity of brain operations. Furthermore, smaller-bodied seed-harvesting species are often sympatric with larger-bodied species and likely compete to some extent for seeds^76^. Though seed sizes are positively correlated with worker body size^83^, interspecific foraging competition^76^ could lead to stronger selection on mass-specific brain performance in smaller-bodied species^41^.

The non-linear relationship between brain MR and brain size provides further evidence that species with smaller-brained workers may be under significant selection to maximise their performance, and therefore brain MRs. Brain MR did not vary between the largest species as much as between the smallest species. The steep increase in brain MR with brain size when brain mass < 0.15 mg indicates that there is a benefit to increasing brain MR at smaller brain sizes, or brain energy consumption is limited at larger brain sizes.

The high brain MSMR of smaller-bodied workers may also be a function of other aspects of miniaturization^47^. Smaller insects preserve axons over glial cells in the central nervous system, likely to maintain brain function. Generating action potentials and synaptic signalling are the costliest neuronal functions^47–50^. Therefore, the necessity to maintain brain function despite miniaturization might require increasing intensity of brain use and high brain MSMRs. Coupled with the strong propensity for sensory organs to also scale hypometrically^101–104^, higher mass-specific information processing for smaller animals could partially preserve cognitive and behavioural capacities^53–56^. This may be a proximate mechanism by which smaller individuals maintain cognitive capacity to compete with larger-bodied sympatric species^41^. Additionally, smaller-bodied species are often more likely to experience predation, which may also select for better sensory systems and reaction times^42^. Further behavioural, ecological, and physiological studies will be required to determine the causal factors driving body-size associated energetic patterns.

Although our metabolic measurements were made on intact undamaged brains, we recognize that the dissection necessary for recording *ex vivo* brain metabolic rates has physiological effects that are not well understood. Severing sensory and motor neurons of the brain will likely first increase ion fluxes and therefore cause apoptosis, though neurons have mechanisms to protect against this process^105^. The number of neurons cut, however, is likely extremely small relative to the total number of brain cells. Smaller *P. rugosus* workers have 50,000-80,000 brain cells and comparable-size workers of other ant species have upwards of 200,000 cells^105^. Furthermore, the stability of brain metabolic rates over several hours demonstrates that dissections are not driving a fast, progressive deterioration of brain function. Data from *in vitro* tissues, including the brain, in fish indicate similar metabolic scaling exponents as those measured from in intact animals^106,107^, and glucose uptake by the brains of starved rats are similarly depressed *in vivo* and *ex vivo*^108^, suggesting that the relationship among metabolic levels of brains are not negatively impacted by removal from the body. Many studies have successfully utilised the Seahorse system to detect the effects of toxins^109^, diet^110^ and trauma, even from partial brains that have undergone destructive sampling^111^. Together these studies suggest that such *ex vivo* brain metabolism measures accurately reflect *in vivo* interspecific patterns, opening avenues for future research.

Size is frequently used as a proxy for brain energetic cost, neuropil investment, and behavioural and/or cognitive processing ability^20^. If absolute brain size is considered as the index of brain investment, one might conclude that the social brain theory was supported in this clade (Fig. 3B); however, incorporation of the effects of size and mass-specific metabolic rate (Figs. 3C, 4C) result in the opposite conclusion. Linkages between brain size (and neuron number) and cognitive ability grow weaker as greater taxonomic diversity is considered^71^, indicating that brain size might be an inconsistent metric for behavioural capability. Our brain MR data provide empirical evidence that brain size is not necessarily an accurate proxy for brain energy use^20,71^. *Pogonomyrmex rugosus* workers have the largest brain in our study (0.229 mg), approximately double the mass of *P. occidentalis* (0.114 mg). However, the difference in whole-brain energy consumption (*P. occidentalis*: 1.76 µW; *P. rugosus*: 1.64 µW) is minimal due to the much higher brain MSMR in the smaller-bodied species. Our study reveals that assessment of brain metabolism in a phylogenetic context can be a powerful tool to understand patterns of brain evolution across diverse animal clades.

## Data and code availability

Data and code used in this manuscript have been deposited at Open Science Foundation and will be publicly available on publication.

## Ethics Statement

The animals used in this study are not subject to any formal legal or ethical oversight. However, care was taken during handling, husbandry and experimentation to minimise any potential suffering.

## METHODS

### i. Study species

Workers of eleven species (Table 1) of *Pogonomyrmex* ants were collected from multiple colonies and locations in Arizona and California, USA between May 2021 and January 2023 (Supplementary Table S1). Ants were housed in small plastic enclosures and stored in an environmental control room with the temperature fixed at 30°C. They were provided with 30% w/v sucrose solution and fresh water *ad libitum* and were processed for experimentation within 3 weeks of collection.

### ii. Sociometrics

We assessed the effect of social organization on brain mass and metabolism using colony size as an estimator of social complexity. Colony size is considered a reliable indicator of derived and advanced sociality in ants^62^ and other eusocial insects^35,52,59,60^ owing to its association with the potential number of interactions among workers, whether or not the relationships among workers or groups of workers are differentiated. Further evidence for colony size as a robust proxy of social complexity can found in *P. barbatus*, where foraging dynamics are modulated by social interaction^112,113.^ The members of these colonies experience high levels of social interaction and form some of the largest colonies in the genus.

### iii. Whole body resting metabolic rate measurements

To determine the whole body metabolic of ants at rest, we used differential flow-through respirometry. The respirometer was in the same temperature-controlled (30°C) room the ants were stored in. We conducted all respirometry trials in the dark, with red filters applied to respirometry chambers to prevent incidental light reaching the ant during preparatory lab work.

A Flowbar-8 mass flow meter system (Sable Systems, Las Vegas, NV, USA) pumped dry, CO_2_ free air from a gas cylinder into the respirometry chamber at a STP flow rate of 50 ± 1 ml min^-1^. Ants were placed into a chamber consisting of Bev-A-Line tube (length: 25mm, diameter: 3.2mm). Output from the chamber was directed to the sample cell of a LI-7000 CO_2_/H_2_O Gas Analyser, which sampled at 1Hz, and was calibrated with two CO_2_ calibration standards analysed ± 0.2 ppm.

We recorded a baseline measurement, without an ant, for at least one minute preceding each respirometry trial. After introducing the ant to the chamber, we covered the chamber with a transparent red filter and allowed the ant to adjust to the chamber until we observed little to no movement. We recorded CO_2_ production and synced this output with ant activity using a web camera (Logitech HD Pro Webcam C920, 1080p) for 30 minutes.

We digitized the analog output from the LI-7000 using a Sable Systems UI2 and recorded once per second using ExpeData (Sable Systems, v. 1.7.2) for Windows. We calculated average CO_2_ levels during time periods when we observed ants to be still.

CO_2_ production rates (ml h^-1^) were calculated using Equation 1, with FR equal to the flow rate (ml h^-1^), and FCO_2_ equal to the fractional CO_2_ level (μmol mol^-1^) in the excurrent air from the respirometry chamber:

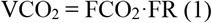

We did not measure the respiratory quotient of all our species. The respiratory quotient of worker ants has been reported as 0.71, 0.77, 0.8, 0.91, 0.92. and 1.02^114–118^. We converted worker CO_2_ production to oxygen consumption using the average respiratory quotient of these studies (0.85) and then calculated metabolic rate in microwatts assuming 20.4 joules ml oxygen^-1 119^. Ants were sacrificed by freezing after being weighed using a XPE56 XPE microanalytical balance (Mettler Toledo, Columbus, OH, USA). Sample sizes for each species can be found in Table S2.

### iv. Brain metabolic rate measurements

Brain oxygen consumption rates were measured *ex vivo* using a Seahorse XF HS Mini Analyser (Agilent, Santa Clara, California, USA) following methods outlined in^120^. Workers were anaesthetised on ice and their fresh masses recorded using an XPE56 XPE microanalytical balance (Mettler Toledo, Columbus, OH, USA). While remaining unconscious from the cold, workers were decapitated. Brains were then removed intact and undamaged from the head capsule under a dissecting microscope in Seahorse XF base media (Agilent, Santa Clara, California, USA) supplemented with 0.01 moles per litre glucose and sodium pyruvate. The Seahorse XF HS Mini Analyser has eight wells, two of which were left empty as control wells, leaving six which contained intact dissected brains. Assays were conducted at 32°C as this was the temperature at which the Seahorse stabilized in the lab with the heater turned off and was very near the holding and respirometry temperature used.

Brain MRs were recorded in cycles. The Seahorse XF HS Mini mixes oxygen into the media within the well and then measures the rate at which oxygen is depleted. This cycle was repeated twelve times, taking approximately an hour; MRs were usually very stable over this period. The rates estimated from the last three cycles were selected and the mean MR was calculated (Fig. S1). The Seahorse XF HS Mini measures oxygen consumption rates (picomols min^-1^). Insect brains are believed to catabolize a mixture of carbohydrates and fatty acid metabolites^121^. In the absence of knowing the exact brain respiratory quotient, we converted oxygen consumption into µwatts using the same conversion factor we used for resting workers, 20.4 joules per ml O_2_ consumed. After completion of the assay, brain mass was measured using a XPE56 XPE microanalytical balance (Mettler Toledo, Columbus, OH, USA). Sample sizes for each species can be found in Table S2.

### v. Statistical analysis

All statistical analyses were conducted using R version 4.2.2^122^. Phylogenetic signal was calculated using the package ‘phytools’^123^, Brownian correlation structures for phylogenetic generalised least squares (PGLS) regressions were calculated using the package ‘ape’^124^ and data visualisation was conducted using the package ‘ggplot2’^125^ and ‘ggtree’^126^. Generalised additive models (GAM) were implemented using the package ‘mgcv’^127^, phylogenetic penalties were implemented using the package ‘MRFtools’^128^.

MRs and brain masses were analysed using linear models constructed using the R base package^122^ or mixed effects models constructed using the ‘lme4’ package^129^. Significance of terms in mixed effects models were assessed using the package ‘lmerTest’^130^. MRs and body masses were log_10_ transformed to facilitate allometric analysis.

### vi. Phylogenetic reconstruction

11 samples from the 143 sample phylogeny inferred in Graber et al.^72^ were used to construct the phylogeny used in our analyses. UCE library prep and enrichment protocols are described in Graber et al ^72^. in prep. 851 alignments with >95% representation of the 11 taxa were concatenated into a single matrix using ‘phyluce_align_get_only_loci_with_min_taxa’. IQ-TREE2 was used to infer a maximum likelihood phylogeny with 1000 bootstrap replicates and the GTR+G model used for the entire alignment.

## REFERENCES

1. Heldstab, S. A., Isler, K., Graber, S. M., Schuppli, C. & van Schaik, C. P. The economics of brain size evolution in vertebrates. Current Biology 32, R697–R708 (2022).

2. Finarelli, J. A. & Flynn, J. J. Brain-size evolution and sociality in Carnivora. Proc Natl Acad Sci U S A 106, 9345–9349 (2009).

3. Chambers, H. R., Heldstab, S. A. & O’Hara, S. J. Why big brains? A comparison of models for both primate and carnivore brain size evolution. PLoS One 16, e0261185 (2021).

4. Watanabe, A., Balanoff, A. M., Gignac, P. M., Gold, M. E. L. & Norell, M. A. Novel neuroanatomical integration and scaling define avian brain shape evolution and development. Elife 10, (2021).

5. Dembitzer, J., Castiglione, S., Raia, P. & Meiri, S. Small brains predisposed Late Quaternary mammals to extinction. Sci Rep 12, (2022).

6. Venditti, C., Baker, J. & Barton, R. A. Co-evolutionary dynamics of mammalian brain and body size. Nature Ecology & Evolution 2024 8:8 8, 1534–1542 (2024).

7. Van Der Woude, E., Smid, H. M., Chittka, L. & Huigens, M. E. Breaking Haller’s Rule: Brain-Body Size Isometry in a Minute Parasitic Wasp. Brain Behav Evol 81, 86–92 (2013).

8. Burger, J. R., George, M. A., Leadbetter, C. & Shaikh, F. The allometry of brain size in mammals. J Mammal 100, 276–283 (2019).

9. Eberhard, W. G. & Wcislo, W. T. Grade changes in brain–body allometry: Morphological and behavioural correlates of brain size in miniature spiders, insects and other invertebrates. in Advances in Insect Physiology (ed. Casas, J.) vol. 40 155–214 (2011).

10. Rensch, B. Histological changes correlated with evolutionary changes of body size. Evolution (N Y) 2, 218–230 (1948).

11. Karbowski, J. Global and regional brain metabolic scaling and its functional consequences. BMC Biol 5, 18 (2007).

12. Dunbar, R. The social brain hypothesis. Evolutionary Anthropology: Issues, News, and Reviews 6, 178–190 (1998).

13. Shultz, S. & Dunbar, R. I. M. Socioecological complexity in primate groups and its cognitive correlates. Philosophical Transactions of the Royal Society B 377, (2022).

14. DeCasien, A. R., Williams, S. A. & Higham, J. P. Primate brain size is predicted by diet but not sociality. Nat Ecol Evol 1, 1–7 (2017).

15. Rosati, A. G. Foraging cognition: Reviving the ecological intelligence hypothesis. Trends Cogn Sci 21, 691–702 (2017).

16. Aiello, L. C. & Wheeler, P. The expensive-tissue hypothesis: The brain and the digestive system in human and primate evolution. Curr Anthropol 36, 199–221 (1995).

17. Navarrete, A., Van Schaik, C. P. & Isler, K. Energetics and the evolution of human brain size. Nature 2011 480:7375 480, 91–93 (2011).

18. Dunbar, R. I. M. & Shultz, S. Why are there so many explanations for primate brain evolution? Philosophical Transactions of the Royal Society B: Biological Sciences 372, (2017).

19. Sobrero, R., May-Collado, L. J., Agnarsson, I. & Hernández, C. E. Expensive brains: ‘Brainy’ rodents have higher metabolic rate. Front Evol Neurosci 3, 10044 (2011).

20. Kverková, K. et al. The evolution of brain neuron numbers in amniotes. Proc Natl Acad Sci U S A 119, e2121624119 (2022).

21. DeSilva, J., Traniello, J., Claxton, A. & Fannin, L. When and why did human brains decrease in size? A new change-point analysis and insights from brain evolution in ants. Front Ecol Evol 9, 742639 (2021).

22. Smaers, J. B. et al. The evolution of mammalian brain size. Sci Adv 7, 9 (2021).

23. Tsuboi, M. et al. Breakdown of brain–body allometry and the encephalization of birds and mammals. Nature Ecology & Evolution 2018 2:9 2, 1492–1500 (2018).

24. Guay, P. J., Weston, M. A., Symonds, M. R. E. & Glover, H. K. Brains and bravery: Little evidence of a relationship between brain size and flightiness in shorebirds. Austral Ecol 38, 516–522 (2013).

25. Nealen, P. M. & Ricklefs, R. E. Early diversification of the avian brain:body relationship. J Zool 253, 391–404 (2001).

26. De Meester, G., Huyghe, K. & Van Damme, R. Brain size, ecology and sociality: A reptilian perspective. Biological Journal of the Linnean Society 126, 381–391 (2019).

27. Taylor, G. M., Nol, E. & Boire, D. Brain regions and encephalization in anurans: Adaptation or stability? Brain Behav Evol 45, 96–109 (1995).

28. Triki, Z., Aellen, M., Van Schaik, C. P. & Bshary, R. Relative brain size and cognitive equivalence in fishes. Brain Behav Evol 96, 124–136 (2021).

29. Quesada, R. et al. The allometry of CNS size and consequences of miniaturization in orb-weaving and cleptoparasitic spiders. Arthropod Struct Dev 40, 521–529 (2011).

30. Polilov, A. A. & Makarova, A. A. The scaling and allometry of organ size associated with miniaturization in insects: A case study for Coleoptera and Hymenoptera. Sci Rep 7, 43095 (2017).

31. Godfrey, R. K., Swartzlander, M. & Gronenberg, W. Allometric analysis of brain cell number in Hymenoptera suggests ant brains diverge from general trends. Proc Biol Sci 288, (2021).

32. Sayol, F. et al. Feeding specialization and longer generation time are associated with relatively larger brains in bees. Proceedings of the Royal Society B 287, (2020).

33. Lösel, P. D. et al. Natural variability in bee brain size and symmetry revealed by micro-CT imaging and deep learning. PLoS Comput Biol 19, e1011529 (2023).

34. Gowda, V. & Gronenberg, W. Brain composition and scaling in social bee species differing in body size. Apidologie 50, 779–792 (2019).

35. Wehner, R., Fukushi, T. & Isler, K. On being small: Brain allometry in ants. Brain Behav Evol 69, 220–228 (2007).

36. Seid, M. A., Castillo, A. & Wcislo, W. T. The allometry of brain miniaturization in ants. Brain Behav Evol 77, 5–13 (2011).

37. Mink, J. W., Blumenschine, R. J. & Adams, D. B. Ratio of central nervous system to body metabolism in vertebrates: Its constancy and functional basis. Am J Physiol 241, 203–212 (1981).

38. Kern, M. J. Metabolic rate of the insect brain in relation to body size and phylogeny. Comparative Biochemistry and Physiology Part A 81, 501–506 (1985).

39. Chittka, L. & Niven, J. Are bigger brains better? Curr Biol 19, R995–R1008 (2009).

40. White, C. R., Alton, L. A., Bywater, C. L., Lombardi, E. J. & Marshall, D. J. Metabolic scaling is the product of life-history optimization. Science (1979) 377, 834–839 (2022).

41. Harrison, J. F. Do performance-safety tradeoffs cause hypometric metabolic scaling in animals? Trends Ecol Evol 32, 653–664 (2017).

42. Glazier, D. S. Does death drive the scaling of life? Biological Reviews 100, 586–619 (2025).

43. Reader, S. M. & Laland, K. N. Social intelligence, innovation, and enhanced brain size in primates. Proceedings of the National Academy of Sciences 99, 4436–4441 (2002).

44. Deaner, R. O., Isler, K., Burkart, J. & van Schaik, C. Overall brain size, and not encephalization quotient, best predicts cognitive ability across non-human primates. Brain Behav Evol 70, 115–124 (2007).

45. Herculano-Houzel, S. Numbers of neurons as biological correlates of cognitive capability. Curr Opin Behav Sci 16, 1–7 (2017).

46. Polilov, A. A. Small is beautiful: features of the smallest insects and limits to miniaturization. Annu Rev Entomol 60, 103–121 (2015).

47. Niven, J. E. & Farris, S. M. Miniaturization of nervous systems and neurons. Current Biology 22, R323–R329 (2012).

48. Sengupta, B., Stemmler, M., Laughlin, S. B. & Niven, J. E. Action potential energy efficiency varies among neuron types in vertebrates and invertebrates. PLoS Comput Biol 6, e1000840 (2010).

49. Attwell, D. & Laughlin, S. An energy budget for signaling in the grey matter of the brain. J Cereb Blood Flow Metab 21, (2001).

50. Quintela-López, T., Shiina, H. & Attwell, D. Neuronal energy use and brain evolution. Current Biology 32, R650–R655 (2022).

51. Farris, S. M. Insect societies and the social brain. Curr Opin Insect Sci 15, 1–8 (2016).

52. O’Donnell, S. et al. Distributed cognition and social brains: Reductions in mushroom body investment accompanied the origins of sociality in wasps (Hymenoptera: Vespidae). Proceedings of the Royal Society B: Biological Sciences 282, 20150791 (2015).

53. Godfrey, R. K. & Gronenberg, W. Brain evolution in social insects: Advocating for the comparative approach. Journal of Comparative Physiology A 2019 205:1 205, 13–32 (2019).

54. Feinerman, O. & Traniello, J. F. A. Social complexity, diet, and brain evolution: modeling the effects of colony size, worker size, brain size, and foraging behavior on colony fitness in ants. Behav Ecol Sociobiol 70, 1063–1074 (2016).

55. Traniello, J. F., Linksvayer, T. A. & Coto, Z. N. Social complexity and brain evolution: insights from ant neuroarchitecture and genomics. Curr Opin Insect Sci 53, 100962 (2022).

56. Coto, Z. N. & Traniello, J. F. A. Social brain energetics: Ergonomic efficiency, neurometabolic scaling, and metabolic polyphenism in ants. Integr Comp Biol 62, 1471–1478 (2022).

57. Lihoreau, M., Latty, T. & Chittka, L. An exploration of the social brain hypothesis in insects. Front Physiol 3, 442 (2012).

58. Traniello, J. F. A. & Avarguès-Weber, A. Individual and collective cognition in social insects: What’s in a name? Behav Ecol Sociobiol 77, 1–12 (2023).

59. Bell-Roberts, L. et al. Larger colony sizes favoured the evolution of more worker castes in ants. Nature Ecology & Evolution 2024 8:10 8, 1959–1971 (2024).

60. Matte, A. & LeBoeuf, A. C. Innovation in ant larval feeding facilitated queen–worker divergence and social complexity. Proceedings of the National Academy of Sciences 122, e2413742122 (2025).

61. Holbrook, C. T., Barden, P. M. & Fewell, J. H. Division of labor increases with colony size in the harvester ant Pogonomyrmex californicus. Behavioral Ecology 22, 960–966 (2011).

62. Anderson, C. & McShea, D. W. Individual versus social complexity, with particular reference to ant colonies. Biol Rev Camb Philos Soc 76, 211–237 (2001).

63. Muratore, I. B., Fandozzi, E. M. & Traniello, J. F. A. Behavioral performance and division of labor influence brain mosaicism in the leafcutter ant Atta cephalotes. Journal of Comparative Physiology A 208, 325–344 (2022).

64. Kamhi, J. F., Gronenberg, W., Robson, S. K. A. & Traniello, J. F. A. Social complexity influences brain investment and neural operation costs in ants. Proceedings of the Royal Society B: Biological Sciences 283, 20161949 (2016).

65. Pahlke, S., Seid, M. A., Jaumann, S. & Smith, A. The loss of sociality is accompanied by reduced neural investment in mushroom body volume in the sweat bee Augochlora pura (Hymenoptera: Halictidae). Ann Entomol Soc Am 114, 637–642 (2020).

66. Smith, A. R., Seid, M. A., Jiménez, L. C. & Wcislo, W. T. Socially induced brain development in a facultatively eusocial sweat bee Megalopta genalis (Halictidae). Proceedings of the Royal Society B: Biological Sciences 277, 2157 (2010).

67. Strausfeld, N. J., Hansen, L., Li, Y., Gomez, R. S. & Ito, K. Evolution, discovery, and interpretations of arthropod mushroom bodies. Learning & Memory 5, 11–37 (1998).

68. Riveros, A. J., Seid, M. A. & Wcislo, W. T. Evolution of brain size in class-based societies of fungus-growing ants (Attini). Anim Behav 83, 1043–1049 (2012).

69. Jerison, H. J. Brain evolution: New light on old principles. Science (1979) 170, 1224–1225 (1970).

70. Cole, A. C. Pogonomyrmex Harvester Ants. A Study of the Genus in North America. (University of Tennessee Press, Knoxville, Tennessee, 1968).

71. Barron, A. B. & Mourmourakis, F. The relationship between cognition and brain size or neuron number. Brain Behav Evol 99, 109–122 (2024).

72. Graber, L. C., Johnson, R. A. & Moreau, C. S. UCE phylogenomics inform the systematics and geographic range evolution of the harvester ant genus Pogonomyrmex. bioRxiv 2024.11.12.623263 (2024) doi:10.1101/2024.11.12.623263.

73. Burger, J. R., Hou, C., A S Hall, C. & Brown, J. H. Universal rules of life: Metabolic rates, biological times and the equal fitness paradigm. Ecol Lett 24, 1262–1281 (2021).

74. Brown, J. H., Burger, J. R., Hou, C. & Hall, C. A. S. The pace of life: Metabolic energy, biological time, and life history. Integr Comp Biol 62, 1479–1491 (2022).

75. Hölldobler, B. Recruitment behavior, home range orientation and territoriality in harvester ants, Pogonomyrmex. Behav Ecol Sociobiol 1, 3–44 (1976).

76. Whitford, W. G. Foraging in seed-harvester ants Pogonomyrmex spp. Ecology 59, 185–189 (1978).

77. Flanagan, T. P., Letendre, K., Burnside, W. R., Fricke, G. M. & Moses, M. E. Quantifying the effect of colony size and food distribution on harvester ant foraging. PLoS One 7, e39427 (2012).

78. Hölldobler, B. Recruitment behavior, home range orientation and territoriality in harvester ants, Pogonomyrmex. Behav Ecol Sociobiol 1, 3–44 (1976).

79. Warburg, I., Whitford, W. G. & Steinberger, Y. Colony size and foraging strategies in desert seed harvester ants. J Arid Environ 145, 18–23 (2017).

80. Weier, J. A. & Jr, D. H. F. Foraging in the seed-harvester ant genus Pogonomyrmex: Are energy costs important? 291–300 (1995).

81. Fewell, J. H. Energetic and time costs of foraging in harvester ants, Pogonomyrmex occidentalis. Behav Ecol Sociobiol 22, 401–408 (1988).

82. Davidson, D. W. Foraging ecology and community organization in desert seed-eating ants. Ecology 58, 725–737 (1977).

83. Davidson, D. W. Species diversity and community organization in desert seed-eating ants. Ecology 58, 711–724 (1977).

84. Wagner, D. & Gordon, D. M. Colony age, neighborhood density and reproductive potential in harvester ants. Oecologia 1999 119:2 119, 175–182 (1999).

85. Waters, J. S., Holbrook, C. T., Fewell, J. H. & Harrison, J. F. Allometric Scaling of Metabolism, Growth, and Activity in Whole Colonies of the Seed-Harvester Ant Pogonomyrmex californicus. Am Nat 176, 501–510 (2010).

86. Fewell, J. H. & Harrison, J. F. Scaling of work and energy use in social insect colonies. Behavioral Ecology and Sociobiology 2016 70:7 70, 1047–1061 (2016).

87. Ostwald, M. M. et al. Cooperation among unrelated ant queens provides persistent growth and survival benefits during colony ontogeny. Sci Rep 11, 8332 (2021).

88. Waters, J. S., Ochs, A., Fewell, J. H. & Harrison, J. F. Differentiating causality and correlation in allometric scaling: Ant colony size drives metabolic hypometry. Proceedings of the Royal Society B 284, 20162582 (2017).

89. Duell, M. E., Klok, C. J., Roubik, D. W. & Harrison, J. F. Size-dependent scaling of stingless bee flight metabolism reveals an energetic benefit to small body size. Integr Comp Biol 62, 1429–1438 (2022).

90. Wagner, J. M. et al. Isometric spiracular scaling in scarab beetles - implications for diffusive and advective oxygen transport. Elife 11, e82129 (2022).

91. Foerster, S. Í. A., Javoiš, J., Holm, S. & Tammaru, T. Predicting insect body masses based on linear measurements: A phylogenetic case study on geometrid moths. Biological Journal of the Linnean Society 141, 71–86 (2024).

92. Herrera, C. M. Thermal biology diversity of bee pollinators: Taxonomic, phylogenetic, and plant community-level correlates. Ecol Monogr 94, e1625 (2024).

93. Taber, S. Welton. The World of the Harvester Ants. (Texas A & M University Press, College Station, 1998).

94. Harrison, J. F. Approaches for testing hypotheses for the hypometric scaling of aerobic metabolic rate in animals. Am J Physiol Regul Integr Comp Physiol 315, R879–R894 (2018).

95. Harrison, J. F., Klok, C. J. & Waters, J. S. Critical PO2 is size-independent in insects: Implications for the metabolic theory of ecology. Curr Opin Insect Sci 4, 54–59 (2014).

96. Burns, J. G., Foucaud, J. & Mery, F. Costs of memory: Lessons from ‘mini’ brains. Proceedings of the Royal Society B: Biological Sciences 278, 923–929 (2011).

97. Shik, J. Z. & Kaspari, M. Lifespan in male ants linked to mating syndrome. Insectes Soc 56, 131–134 (2009).

98. Turza, F., Stec, D., Fontaneto, D. & Miler, K. Life expectancy in ants explains variation in helpfulness regardless of phylogenetic relatedness. Behavioral Ecology https://doi.org/10.1093/beheco/arae104 (2024) doi:10.1093/beheco/arae104.

99. Koch, R. E. et al. Integrating mitochondrial aerobic metabolism into ecology and evolution. Trends Ecol Evol 36, 321–332 (2021).

100. Shilovsky, G. A., Putyatina, T. S. & Markov, A. V. Evolution of longevity in tetrapods: Safety is more important than metabolism level. Biochemistry (Moscow) 89, 322–340 (2024).

101. Perl, C. D. & Niven, J. E. Colony-level differences in the scaling rules governing wood ant compound eye structure. Sci Rep 6, 24204 (2016).

102. Chong, K. L., Grahn, A., Perl, C. D. & Sumner-Rooney, L. Allometry and ecology shape eye size evolution in spiders. Current Biology 34, 3178–3188.e5 (2024).

103. Brooke, M. D. L., Hanley, S. & Laughlin, S. B. The scaling of eye size with body mass in birds. Proceedings of the Royal Society of London B: Biological Sciences 226, 405–412 (1999).

104. Huang, C. H., Zhong, M. J., Liao, W. B. & Kotrschal, A. Investigating the role of body size, ecology, and behavior in anuran eye size evolution. Evol Ecol 33, 585–598 (2019).

105. Aydin, M. S. et al. Active shrinkage protects neurons following axonal transection. iScience 26, 107715 (2023).

106. Oikawa, S. & Itazawa Yasuo. Relationship between summated tissue respiration and body size in a marine teleost, the porgy Pagrus major. Fisheries Science 69, 687–694 (2003).

107. Oikawa, S. & Itazawa, Y. Relationship between metabolic rate in vitro and body mass in a marine teleost, porgy Pagrus major. Fish Physiol Biochem 10, 177–182 (1992).

108. Buschiazzo, A. et al. Effect of starvation on brain glucose metabolism and 18F-2-fluoro-2-deoxyglucose uptake: An experimental in-vivo and ex-vivo study. EJNMMI Res 8, (2018).

109. Moncada-Restrepo, M. & Chambers, J. W. Respirometry parameters as indicators of neurotoxicity. Adv Neurotoxicol 13, 55–80 (2025).

110. Mackert, O. et al. Impact of metabolic stress induced by diets, aging and fasting on tissue oxygen consumption. Mol Metab 64, (2022).

111. Underwood, E., Redell, J. B., Zhao, J., Moore, A. N. & Dash, P. K. A method for assessing tissue respiration in anatomically defined brain regions. Scientific Reports 2020 10:1 10, 1–14 (2020).

112. Gordon, D. M. Chapter 4: Colony Size. in Ant Encounters: Interaction Networks and Colony Behavior 75–95 (Princeton University Press, 2010).

113. Gordon, D. M. The rewards of restraint in the collective regulation of foraging by harvester ant colonies. Nature 498, (2013).

114. Nielsen, M. G., Jensen, T. F. & Holm-Jensen, I. Effect of load carriage on the respiratory metabolism of running worker ants of Camponotus herculeanus (Formicidae). Oikos 39, 137 (1982).

115. Duncan, F. D. & Lighton, J. R. B. The burden within: The energy cost of load carriage in the honeypot ant, Myrmecocystus. Physiol Zool 67, 190–203 (1994).

116. Vogt, J. T. & Appel, A. G. Standard metabolic rate of the fire ant, Solenopsis invicta Buren: effects of temperature, mass, and caste. J Insect Physiol 45, 655–666 (1999).

117. Nielsen, M. G. & Christian, K. A. The mangrove ant, Camponotus anderseni, switches to anaerobic respiration in response to elevated CO2 levels. J Insect Physiol 53, 505–508 (2007).

118. Baroni-Urbani, C. & Nielsen, M. G. Energetics and foraging behaviour of the European seed harvesting ant Messor capitatus: II. Do ants optimize their harvesting? Physiol Entomol 15, 449–461 (1990).

119. Lighton, J. R. B. Measuring Metabolic Rates: A Manual for Scientists. (Oxford University Press, 2019).

120. Neville, K. E. et al. A novel ex vivo method for measuring whole brain metabolism in model systems. J Neurosci Methods 296, 32–43 (2018).

121. Rittschof, C. C. & Schirmeier, S. Insect models of central nervous system energy metabolism and its links to behavior. Glia 66, 1160–1175 (2018).

122. R Core Team. R: A language and environment for statistical computing. Preprint at https://www.r-project.org (2024).

123. Revell, L. J. phytools: an R package for phylogenetic comparative biology (and other things). Methods Ecol Evol 3, 217–223 (2012).

124. Paradis, E. & Schliep, K. ape 5.0: An environment for modern phylogenetics and evolutionary analyses in R. Bioinformatics 35, 526–528 (2019).

125. Wickham, H. Ggplot2: Elegant Graphics for Data Analysis. (Springer-Verlag New York, 2016).

126. Yu, G. Using ggtree to visualize data on tree-like structures. Curr Protoc Bioinformatics 69, e96 (2020).

127. Wood, S. N. Fast stable restricted maximum likelihood and marginal likelihood estimation of semiparametric generalized linear models. J R Stat Soc Series B Stat Methodol 73, 3–36 (2011).

128. Pedersen, E. & Simpson, G. MRFtools: Tools for constructing and plotting Markov Random Fields in R for Graphical Data. R package version 0. 0–2, commit 57dc21b8d46050af443d5a3c9d397e425f09914b. https://github.com/eric-pedersen/MRFtools (2023).

129. Bates, D., Mächler, M., Bolker, B. M. & Walker, S. C. Fitting linear mixed-effects models using lme4. J Stat Softw 67, 1–48 (2015).

130. Kuznetsova, A., Brockhoff, P. B. & Christensen, R. H. B. lmerTest package: Tests in linear mixed effects models. J Stat Softw 82, 1–26 (2017).

131. Cole, A. C. Studies of New Mexico ants. IX. Pogonomyrmex apache (Wheeler) a synonym of Pogonomyrmex sancti-hyacinthi (Wheeler) (Hymenoptera: Formicidae). Journal of the Tennessee Academy of Sciences 29, 266–271 (1954).

132. Tschinkel, W. R. Lifespan, age, size-specific mortality and dispersion of colonies of the Florida harvester ant, Pogonomyrmex badius. Insectes Soc 64, 285–296 (2017).

133. Gordon, D. M. Nest relocation in harvester ants. Ann Entomol Soc Am 85, 44–47 (1992).

134. Creighton, W. S. Studies on the North American representatives of Ephebomyrmex (Hymenoptera: Formicidae). Psyche (Camb Mass) 63, 54–66 (1956).

135. Creighton, W. S. Studies on Arizona ants (3) the habits of Pogonomyrmex huachucanus (Wheeler) and a description of the sexual castes. Psyche (Camb Mass) 59, 71–81 (1952).

136. Heinze, J., Hölldobler, B. & Cover, S. P. Queen polymorphism in the North American harvester ant, Ephebomyrmex imberbiculus. Insectes Soc 39, 267–273 (1992).

137. Lavigne, R. J. Bionomics and nest structure of Pogonomyrmex occidentalis (Hymenoptera: Formicidae). Ann Entomol Soc Am 62, 1166–1175 (1969).

138. Johnson, R. A., Holbrook, C. T., Strehl, C. & Gadau, J. Population and colony structure and morphometrics in the queen dimorphic harvester ant, Pogonomyrmex pima. Insectes Soc 54, 77–86 (2007).

139. MacKay, W. P. A comparison of the energy budgets of three species of Pogonomyrmex harvester ants (Hymenoptera: Formicidae). Oecologia 66, 484–494 (1985).

