## Supplemental Information for "Evolution of neurometabolic frugality in harvester ants"

SUPPLEMENTARY MATERIAL

*Table S1: Sample sizes from each species*

| Species | Metabolic rate type | Number colonies | Number individuals |
|---|---|---|---|
| *P. apache* | Body | 3 | 15 |
| | Brain | 2 | 11 |
| *P. badius* | Body | 3 | 19 |
| | Brain | 3 | 19 |
| *P. barbatus* | Body | 10 | 51 |
| | Brain | 1 | 17 |
| *P. californicus* | Body | 3 | 13 |
| | Brain | 3 | 15 |
| *P. desertorum* | Body | 2 | 36 |
| | Brain | 3 | 15 |
| *P. huachucanus* | Body | 3 | 26 |
| | Brain | 3 | 7 |
| *P. imberbiculus* | Body | 2 | 12 |
| | Brain | 3 | 13 |
| *P. maricopa* | Body | 3 | 16 |
| | Brain | 2 | 10 |
| *P. occidentalis* | Body | 4 | 14 |
| | Brain | 3 | 17 |
| *P. pima* | Body | 3 | 36 |
| | Brain | 3 | 5 |
| *P. rugosus* | Body | 2 | 14 |
| | Brain | 1 | 7 |

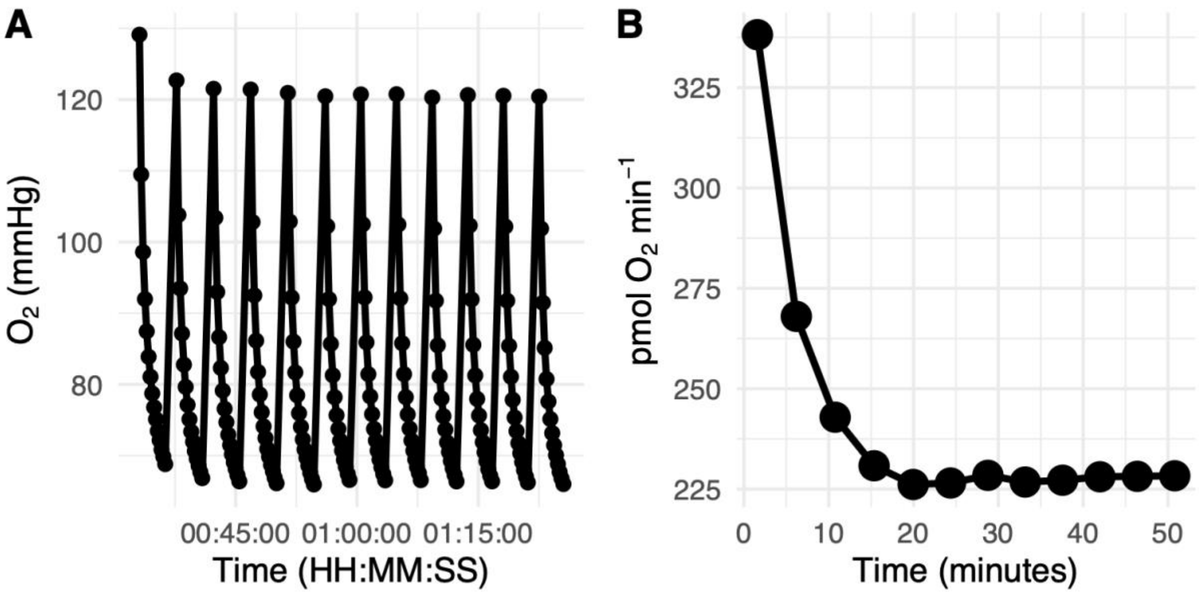


**Fig. S1:** Representative brain metabolic rate data from Agilent Seahorse.

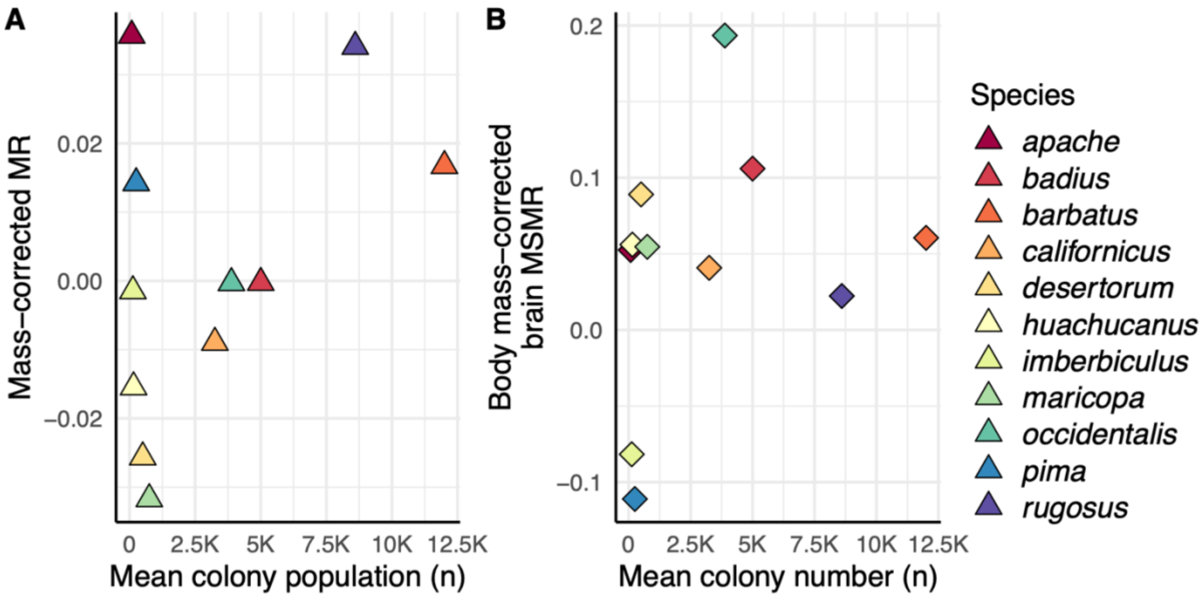


**Fig. S2: Body mass corrected metabolic rates as a response to colony size. A)** Body mass-corrected whole-body metabolic rate (residuals of whole-body MR regressed against body mass, Fig. 4A) as a response to mean colony population. **B)** body-mass-corrected brain MSMR (residuals of brain MSMR regressed against body mass, Fig. 5D) as a response to mean colony population.
